# GENPPI: standalone software for creating protein interaction networks from genomes

**DOI:** 10.1101/2021.01.10.426094

**Authors:** William Ferreira, Gabriel Lanes, Vasco Azevedo, Anderson Santos

## Abstract

**Motivation:** Bacterial genomes are being deposited into online databases at an increasing rate. Genome annotation represents one of the first efforts to understand organisms and their diseases. Some evolutionary relationships that are capable of being annotated only from genomes are conserved gene neighbourhoods (CNs), phylogenetic profiles (PPs), and gene fusions. At present, there is no standalone software that enables networks of interactions among proteins to be created using these three evolutionary characteristics with efficient and effective results.

**Results:** We developed GENPPI software for the *ab initio* prediction of interaction networks using predicted proteins from a genome. In our case study, we employed 50 genomes of the genus *Corynebacterium*. Based on the PP relationship, GENPPI differentiated genomes between the ovis and equi biovars of the species *Corynebacterium pseudotuberculosis* and created groups among the other species analysed. If we inspected only the CN relationship, we could not entirely separate biovars, only species. Our software GENPPI was determined to be efficient because, for example, it creates interaction networks from the central genomes of 50 species/lineages with an average size of 2200 genes in less than 40 minutes on a conventional computer. Our software is compelling because the interaction networks that it creates reflect evolutionary relationships among species and were obtained in average nucleotide identity (ANI) analyses. Additionally, this software enables the user to define how he or she intends to explore the PP and CN characteristics through various parameters, enabling the creation of customized interaction networks. For instance, users can set parameters regarding the genus, metagenome, or pangenome. In addition to the parameterization of GENPPI, it is also the user’s choice regarding which set of genomes he or she is going to study.

**Availability:** The source code in the Common Lisp language, binary files for different operating systems, and GENPPI software tutorials are available at {{github.com/santosardr/genppi}}.

**Supplementary information:** Supplementary data are available at *Bioinformatics* online.

## 1 Introduction

Producing an interaction network is relatively simple; researchers simply need to find a reason to link pairs of entities and apply this rule for all possible pairs of a set. However, such a reason should be trustworthy, or else we could have messy, random, and ineffective relationships. Considerable time and resources could be lost in explaining a non-existent solution for an annotated relation among subjects. Thus, the fundamental role of always present and useful databases becomes clear. Some notable data sources for genome annotation include the following:

- Search Tool for the Retrieval of Interacting Genes/Proteins (STRING) (Szklarczyk *et al.*, 2019);
- Database for Annotation, Visualization and Integrated Discovery (DAVID) (Jiao *et al.*, 2012);
- Metascape (Zhou *et al.*, 2019);
- Kyoto Encyclopedia of Genes and Genomes (KEGG) (Kanehisa *et al.*, 2018);
- Gene Ontology (GO) (“The Gene Ontology Resource: 20 Years and Still GOing Strong”, 2018); and
- Gene Expression Omnibus (Clough and Barrett, 2016).

These well-known databases possess easy-to-use enrichment analyses and useful and user-friendly interfaces for biologists. Many of these databases allow researchers to export their results and continue additional studies using various programs, such as Python (Van Rossum and Drake, 2009), Cytoscape (Shannon, 2003), R (R Core Team, 2013), UALCAN (Chandrashekar *et al.*, 2017), MCODE (Bader and Hogue, 2003), and GEPHI (Leonard, 2004). Notably, there are a considerable number of libraries existing and deployed annually for all this software. For instance, such libraries enable researchers to focus on candidate hub genes, differentially expressed genes (DEGs), the tertiary structure of protein interactions, and many other useful features. For example, in (Sun and Zhang, 2020), the authors studied crucial genes in hepatocellular cancer. The authors obtained the initial data from the Gene Expression Omnibus database. The DAVID website was employed to perform the GO and KEGG enrichment analyses before uploading the data to the STRING database, which was utilized for further analysing the DEGs. After that step, the authors used Cytoscape software to construct a protein interaction network. Once in Cytoscape, a plugin for MCODE was used to study the modules of DEGs. For a final analysis, the authors used the Gene Expression Profiling Interactive Analysis website to determine the module genes’ effects on overall survival under hepatocellular cancer. (Sun and Zhang, 2020) employs a notably elaborate combination of several databases and software tools to produce interesting in silico bioinformatic analyses. There are many other studies similar to this one. A simple internet search of the main terms in this section could retrieve dozens of thousands of similar studies.

Many of the cited databases in this section have the common characteristic of being sealed databases. We define sealed as not accepting new data from anyone outside a trained and specialized team of workers. There is nothing wrong with this approach; one does not allow others to access their bank accounts because of such concerns regarding unauthorized access. For instance, one cannot upload a new genome to the STRING database. First, the database administrator must ensure that the data are trustworthy. Second, a new genome should have some representativeness level to acquire a specific matching of annotation according to the genomes already in the database to reduce the risk of producing poor annotations. Nonetheless, many users would prefer to have their novel genomes annotated by such useful software. Indeed, users can upload their novel genomes to the STRING database and subject them to various kinds of enrichment according to a plethora of third-party databases but only for known genes, not for novel genes. A researcher investigating model organisms will not face such challenges in obtaining useful insights from all the databases mentioned earlier.

For instance, when studying *H. sapiens, M. musculus, R. norvegicus, D. rerio, D. melanogaster, C. elegans, S. cerevisiae, A. thaliana, S. pombe,* and *P. falciparum*, if the STRING (Szklarczyk *et al.*, 2019), Metascape (Zhou *et al.*, 2019), and DAVID (Jiao *et al.*, 2012) databases are employed, a list of genes is sufficient to provide useful data. However, when investigating unseen or underrepresented organisms, a researcher will not have a trustworthy list of genes. Many of the open reading frames (ORFs) will be of unknown function. Such a scenario is more likely to occur when studying prokaryotes. The study of prokaryotes yields dozens of novel genomes and thousands of novel genes daily. We believe that these novel data, even those not curated, deserve the benefit of doubt and further annotation, including topological annotations. We are also confident that the currently utilized databases will not easily manage such a massive volume of novel data. We support the parallelism of this considerable data novelty processing by the creators of the data, the researchers, not by centralized databases, at least in the early stages of data generation. To achieve our vision of parallelism, we developed GENPPI software. GENPPI transfers the question of topological annotation from the centralized databases to the final user, the researcher, at the initial point of research. GENPPI enables researchers to experiment among better sets of genomes to create topological annotation. For instance, we believe that the GENPPI topological annotation information is directly proportional to the number of genomes used to create an annotation. In contrast, the data are indirectly proportional to the number of genomes used for a GENPPI round. As we employ fewer genomes in an annotation round, GENPPI will suggest more interactions between the ORFs, since there are not too many genomes to confirm such a set of predictions as co-occurring. We constantly search for equity between data and information but are guided by the skills of researchers regarding the organisms under study.

GENPPI inspects genomes represented as proteins in the multifasta format, searching for a conserved neighbourhood, phylogenetic profile, and gene fusion. This software enables the decision of how many and what genomes to use for the construction of a protein interaction network to be transferred to the final user. Despite the limited number of features employed in GENPPI, in the next sections, we will attempt to demonstrate that this set of characteristics suffices to produce good-quality networks. We will attempt to support our hypotheses based on the construction of finely detailed phylogenetic maps for the genomes under study. We will demonstrate that the features used by GENPPI can distinguish between, for example, the biovars of the species *Corynebacterium pseudotuberculosis* (Bernardes *et al.*, 2020), as well as obtaining optimal separation among the genera of other prokaryotic organisms, although the software is not limited to one-cell organisms. Considering the quality of species separation and based on the three features analysed by GENPPI, our software obtained good quality for our topological annotations, as well as fewer computational resources needed for this task. For instance, for 50 genomes of an organism containing an of average 2200 genes, we spent only a matter of hours accomplishing full topological annotation.

## 2 Methods

### 2.1 Genome studied

We obtained the genomes investigated in this work using the official NCBI file transport protocol (Supplementary Method S1).

### 2.2 Metrics and reference genomes

To test the validity of the results observed within the GENPPI interaction networks, we performed trials with variations in the following parameters. We describe metrics 1 to 5 as the following:

1. Number of nodes/vertices: number of proteins present in the network;
2. Average degree: number of existing interactions compared to the number of proteins;
3. Density: ratio between a total number of edges and possible edges according to the number of vertices;
4. Number of edges: number of interactions between the proteins in the network;
5. Maximum degree: number of interactions that the most interactive protein has within the network.

We obtained the interaction networks of a set of genomes from model organisms from the STRING database. We calculated these metrics using the software GEPHI and ordered the columns according the level of importance (Supplementary Table S2).

### 2.3 Novel heuristic for faster sequence proteins comparing

In our software GENPPI, we represent the proteins through an amino acid histogram, which indicates the amino acid frequency distribution within a protein sequence. In the process of comparing two proteins, we applied our similarity heuristic approach, known as Histofasta checking (Supplementary Method S3).

## 3 Results

Fig. 1 presents a scheme in which we attempt to explain the disposition of the results that we obtained with GENPPI. Since the GENPPI program can show neighbourhood conservation or phylogenetic profiles, the first step is to produce a pangenome. The data on this pangenome are not in this session of results but rather are results derived from the pangenome. In possession of a pangenome, GENPPI conducts a systematic search for neighbourhoods and conserved phylogenetic profiles. To direct this search, we start from proteins with a high identity (> 90%) or proteins with a high chance of belonging to a central or accessory genome under analysis. The point of this approach is to show that the central pangenome’s characteristics, phylogenetic profile (PP), and conserved neighbourhood (CN) are reliable; otherwise, they would not correctly represent facts about the evolutionary relationship of known bacterial species. Once the correction of evolutionary relationships is confirmed, we can explore distinct ways of generating these networks. In brief, the network creation process variations stem from limitations that we can attribute to how many interactions we want to be part of formatted networks to answer a specific scientific question. However, regardless of the level of data restriction imposed by the user to answer their scientific query, we ensure that the networks produced by GENPPI are reliable because they represent, with a high degree of confidence, the evolutionary relationships of the bacterial species under analysis.

**Fig. 1:**
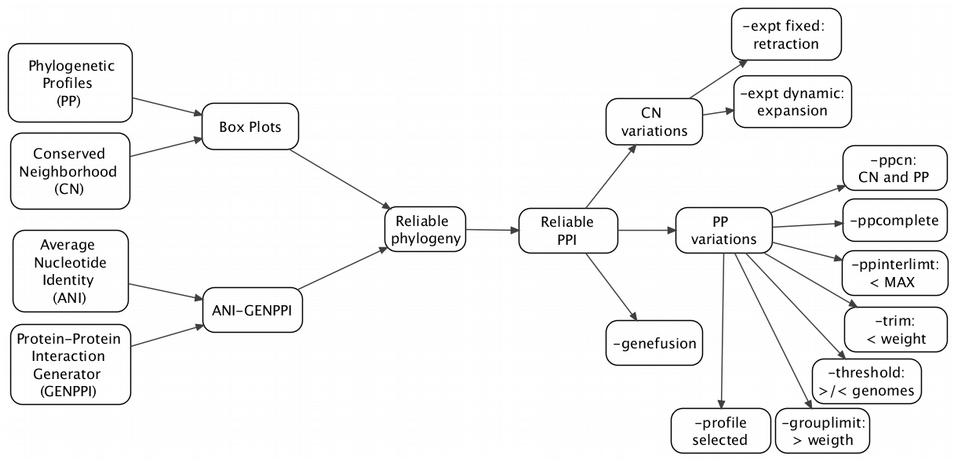
Scheme of an arrangement of the results. Suggesting that GENPPI produces reliable results.

### 3.1 Heat-Maps

The analysis of the difference between genomes using nucleotide sequences, known as Average Nucleotide Identity (ANI), is presented in Figures. Fig. 3 depict the results of GENPPI for the same genomes. However, the data used in Fig. 3 show the extent of proteins shared between each pair of genomes. For example, suppose genome A has 2200 proteins. Of this total, 2000 proteins of genome A have high similarity to proteins of genome B. Therefore, at row A and column B of the heat graph, we have 2000/2200=0.91% protein similarity between genomes A and B. Note that in row B and column A, the protein similarity value between these genomes is likely to be specific. We explain this difference as occurring because the denominator is the measure of B proteins, and the numerator is the chunk of B proteins found in A. The cell colours above and below the main diagonal depend on which genome is the numerator and which is denominator. In Figures 2 and 3, we chose the colours white and black for low and high identical genomes, respectively. The gray colour is an intermediate value between white and black.

**Fig. 2:**
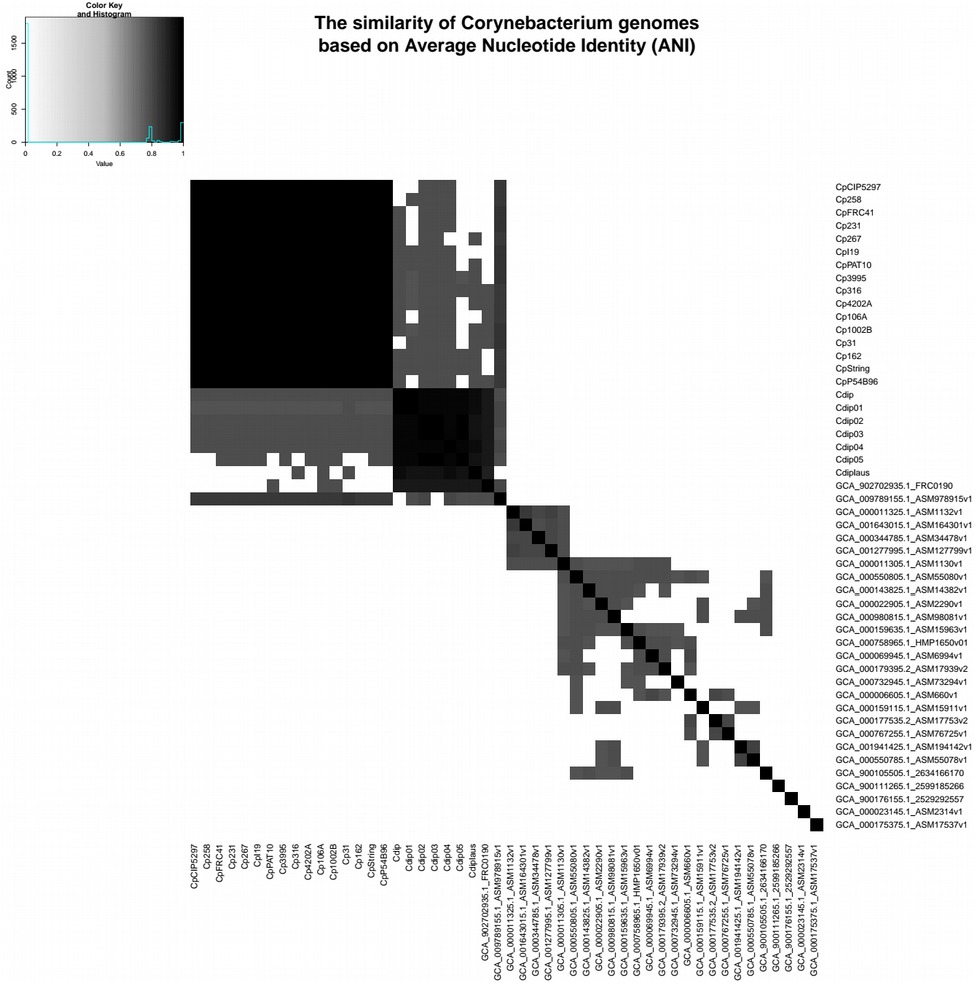
Average nucleotide identity for 50 genomes of the genus *Corynebacterium*. The largest grayish square represents the clusters of *C. pseudotuberculosis* and *Corynebacterium diphtheriae*. Both groupings have units that are almost black due to the high score of DNA similarities.

**Fig. 3:**
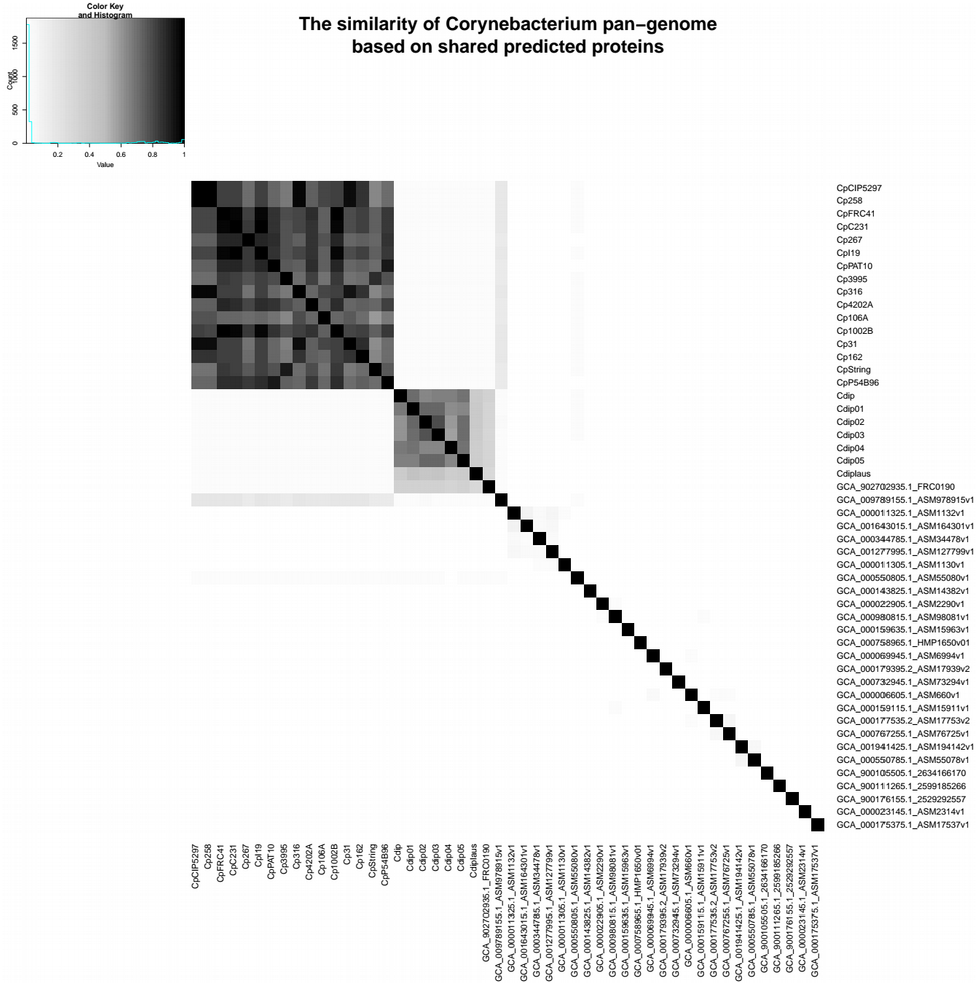
Pangenome similarity profile for the same 50 genomes of the genus *Corynebacterium* depicted in Fig. 2. The clusters of *C. pseudotuberculosis* and *Corynebacterium diphtheriae* are the most grayish. The remaining units are whitish due to the low protein similarities of their phylogenetic profiles.

Genomes of correlated species compared by ANI are differentiated by small percentages and are generally above 90% (Fig. 2). Values of protein similarity between the pangenome (Fig. 3) were less sharpened than the ANI values. A rate of less than 50% can be a high similarity value between a pair of genomes. Supplementary Result Fig. 1 displays the cumulative probability distribution function or CDF plot for the similarities in Fig. 3. The majority (87%) of the possible combinations obtained from the 50 genomes of the genus *Corynebacterium* have a similarity of less than 50%.

Fig. 4 represents the differences between the similarities of each pair of genomes, as determined by ANI (Fig. 2) - GENPPI (Fig. 3). Importantly, the differences indicated in Fig. 4 are not regarding the similarity between the species but how much GENPPI and ANI on these species agree or diverge. In Fig. 4, heat map cells with black values indicate a very pronounced difference, while white values indicate a slightly significant difference between ANI and GENPPI. Most of the units that constitute the *C. pseudotuberculosis* grouping are white. Other units are slightly grayish, representing differences with little expressiveness, between ANI and GENPPI. Excluding the Cdiplaus genome, the *Corynebacterium diphtheriae* cluster would have a colour pattern similar to that of *C. pseudotuberculosis*.

**Fig. 4:**
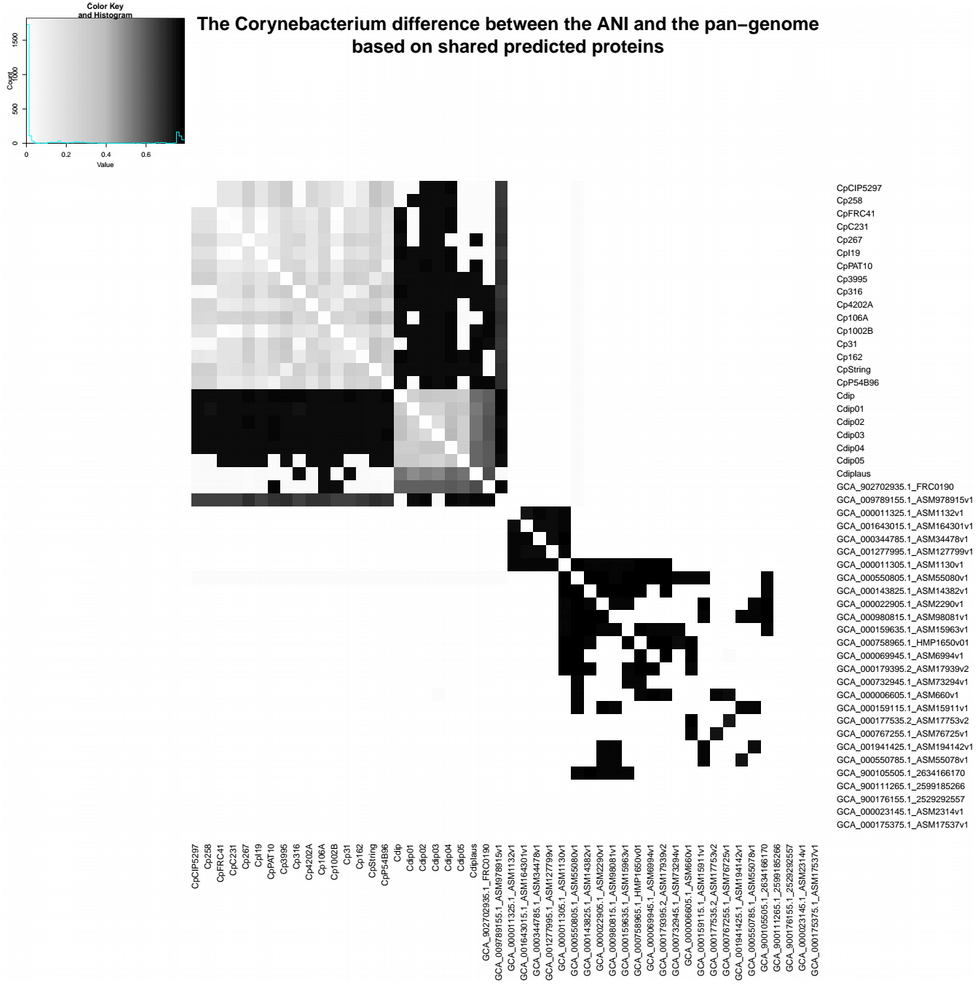
Profile differences between Figures 2 and 3. It accounts for the chunk of divergence about ANI and the pangenome raised by GENPPI. Black cells represent the maximum difference, while white cells account for smaller differences, with grayish units representing intermediate differences. The majority of the comparisons are white because ANI and GENPPI agree on the low similarity of the compared genomes and small differences.

Some cases are noteworthy in Figures 2–4. (i) The genome identified as GCA_902702935.1_FRC0190 refers to *Corynebacterium rouxii* (high GC Gram+). This genome showed high similarity at both the nucleotide and protein levels with the *C. diphtheriae* grouping. An analysis of the data from the heat maps of our work indicates that the genome named *C. rouxii* was *C. diphtheriae*. In addition to our analyses, the specialized literature in these organisms confirms our recommendation to change the nomenclature from the species *C. rouxii* to *C. diphtheriae* (Badell *et al.*, 2020). (ii) The genome identified as GCA_009789155.1_ASM978915v1 refers to *Corynebacterium ulcerans*, strain MRi49. According to the ANI analysis, this genome exhibited high similarity at the nucleotide level with the clusters of *C. pseudotuberculosis* and *C. diphtheriae*. However, the genome exhibited a higher similarity at the protein level with *C. pseudotuberculosis*. Nevertheless, given that we can perceive a slight gray colour in the GENPPI heat map, we believe that this species has some protein similarity to *C. pseudotuberculosis*. In this case, the literature describes the species *C. pseudotuberculosis*, *C. diphtheriae*, and *C. ulcerans* as being evolutionarily related (Busch *et al.*, 2019) (McNamara, Cuevas, and Songer 1995).

Most of Fig. 4 is coloured white, meaning that the ANI enables us to reach the same conclusion as GENPPI regarding the minor similarity between the majority of the possible relationships between each pair of genomes. However, there is a considerable portion of Fig. 4 that is in black colouration. The colour reflects similarities found at the nucleotide level that do not sustain themselves at the amino acid level compared with the pangenome analyses of GENPPI. It is interesting to note that for the clusters of *C. pseudotuberculosis* and diphtheriae, the pattern of similarity between ANI and GENPPI is notable, despite the presence of other numerical values. By guarding the differences in the similarity quantities, we reach the same conclusions between Figures 2 and 3 regarding the evolutionary proximity of cluster organisms. Using the ANI results (Fig. 2), we can note similarities between genome sequences not reflected in the pangenome (Fig. 3). Such closeness extends beyond the clusters of *C. diphtheriae* and *C. pseudotuberculosis*. Therefore, our results support the hypothesis that the similarity between species using the protein pangenome is more useful for differentiating them compared to the DNA sequences. This finding is reasonable because we have demonstrated in Fig. 3 that the species are distinctive with distinct pangenomes, despite having similar DNA sequences, as depicted in Fig. 2. The differentiation of proteins is known to occur because of transcription in the DNA strands. Therefore, a phylogenetic analysis using the pangenome helps to more accurately determine differences between species compared with an identical study examining DNA. Nevertheless, when we analysed genomes from the same species, there was parity between phylogenetic analyses using ANI and GENPPI.

### 3.2 Graph of boxes of conserved phylogenetic profiles

Fig. 5 summarizes the phylogenetic profiles present in each genome analysed (Genome) versus the bulk of genomes in which these profiles appear (Genomes). Therefore, the Y-axis is on the scale from 0 to N, where N is the total genome. In this graph, a median means a load of genomes in which we found conserved phylogenetic profiles, and the width of a plot box is proportional to the number of profiles conserved in a genome. The analysis of conserved phylogenetic profiles made by GENPPI demonstrated the relationship between the ovis and equi biovars of *C. pseudotuberculosis*. The biovar equilinage has six genomes: Cp106A, Cp162, Cp258, Cp31, Cp316, and CpCIP5297. We employed the median and first and third quartiles of genome box plots to demonstrate biovar equi separation. The equi biovar is represented by the first quartile of the plot boxes of the genomes aligning near the median of the plot boxes of genomes belonging to the biovar ovis. The genomes of the biovar equi that fall into this scenario are the following: GCA_000265545.3_ASM26554v3 (Cp162, from a camel in Egypt), GCA_000263755.3_ASM26375v3 (Cp258, from a horse), GCA_000259155.4_ASM25915v4 (CpCp 31, from a buffalo), GCA_000248375.2_ASM24837v2 (Cp316, from a horse in the USA) and GCA_000227605.3_ASM22760v3 (CpCIP5297, from a horse in Kenya). The exception to this rule was the genome with end GCA_000233735.1_ASM23373v1 (Cp106A, from a horse in the USA), which presented the first quartile closest to the ovis biovar strains. However, the median Cp106A was observed to be closer to the biovar equi.

**Fig. 5:**
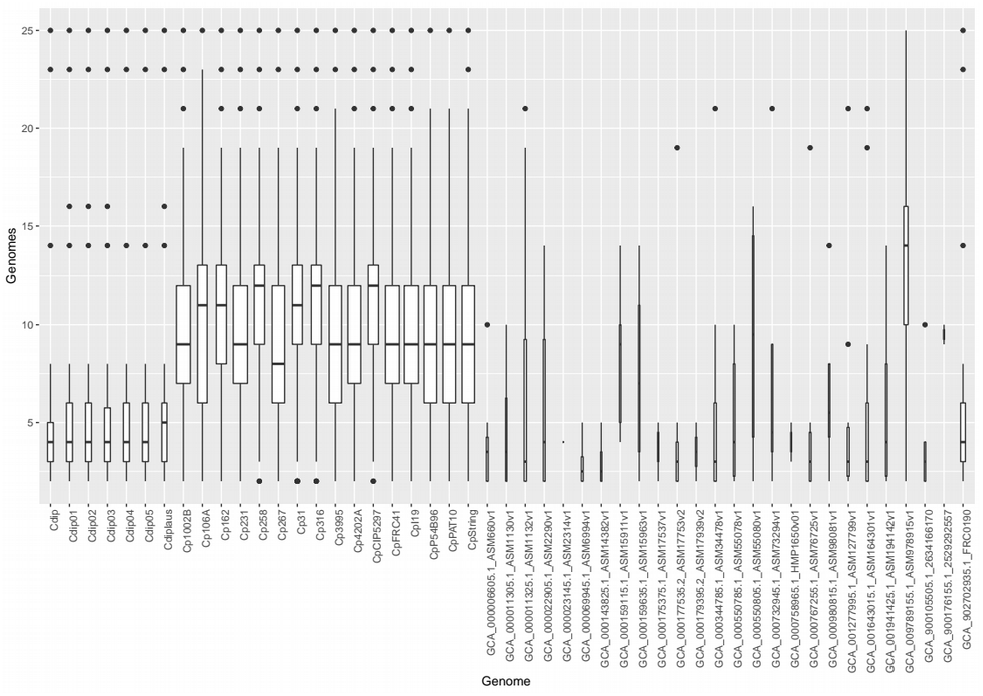
Each genome has a box plot registering stats for their conserved phylogenetic profiles. The width of a box plot is proportional to the number of PPs found. There is a numerical expressiveness of genomes from *C. pseudotuberculosis* (left) and *diphtheriae* (leftmost). For the former, the PP median enables separation of the biovars ovis and equi (highest medians).

The expanded box plots are sixteen and comprise the species *C. pseudotuberculosis*. The box plots of the genomes of the species *C. diphtheriae* are seven and have a smaller width than that of *C. pseudotuberculosis*. Even because these box plots are less represented in this set of genomes, the other species did not show expressive phylogenetic conservation, and we presented plot boxes with a small width. These other species have only one genome representing them in this set of 50 from the genus *Corynebacterium*. Genomes numerically underrepresented compared to *C. pseudotuberculosis* and *C. diphtheriae* account for phylogenetic profiles preserved solely for the genus *Corynebacterium*. The *C. diphtheriae* and *C. pseudotuberculosis* clusters, on the other hand, dominate the number of conserved phylogenetic profiles.

We utilized the species *C. diphtheriae* as a reference genome to assemble the first fifteen genomes of *C. pseudotuberculosis*. At the time, we believed that the species *C. diphtheriae* and *C. pseudotuberculosis* were very similar. At the end of the first assembly, we concluded that these species had a similarity level above 60% at the protein level. For the first automatic annotation transfer, this level of similarity was satisfactory. However, in Fig. 3, the colouration of protein similarity between Cp1002 and Cdip can be observed to be intense white staining, which reflects 2.4% protein similarity with a confidence level greater than 90% of the pangenome. This similarity is low because we set the program to raise the pangenome between these two strains to consider proteins similar only if they had more than 90% identity at the amino acid level. If we had decreased the criterion for determining resemblance, there would probably be a greater affinity between these two species. However, if we had diminished the stringency for proteins’ identity to nearby levels, 60% GENPPI would not translate such a set, given the pangenome’s reliability. With low levels of similarity, preserved protein domains that are present in many proteins with distinct functions could lead to false positive results regarding the pangenome’s central genome.

The previous section showed the utility of generating a central genome with the ability to create phylogenetic clusters consistent with our biological knowledge of bacterial species. The analysis of the box chart results in Fig. 5 shows that phylogenetic profiles made by GENPPI are also consistent with the previous findings regarding species and biovars. Thus, the interaction networks created by GENPPI using the conservation of phylogenetic profiles can help us to identify a topological structure with biological significance.

### 3.3 Box plot of preserved gene neighbourhoods

GENPPI does not work with the genomic DNA sequence but with a report exported from the DNA encoding proteins. However, the conservation of a gene’s DNA sequence location influences the box plot of preserved gene neighbourhoods. We assume that protein sequences tend to enter a multifasta file in an order similar to that observed when they were when extracted from a DNA sequence. GENPPI software receives as input a multifasta file of proteins ordered similar to the corresponding genes arranged on the DNA sequence. Given this premise, in Fig. 6, we use a window of size w to count how many genes are conserved according to at least some other N genomes under analysis. We store a conservation pattern if that pattern occurs in two or more genomes. Two very similar genomes may have almost identical gene neighbourhoods. For an example of two genomes evolutionarily close and assuming a value of w < 10, the median of a conserved neighbourhood (CN), the first quartile and the third quartile, as well as the maximum number of conserved genes, are all equal to w, except for several outliers. The greater the extent of a box plot is, the greater the number of genes with CN characteristics in a genome is. In a CN graph, there is no way to know which genomes are very similar. It is possible to know that there are very similar genomes with a minimum of two.

**Fig. 6:**
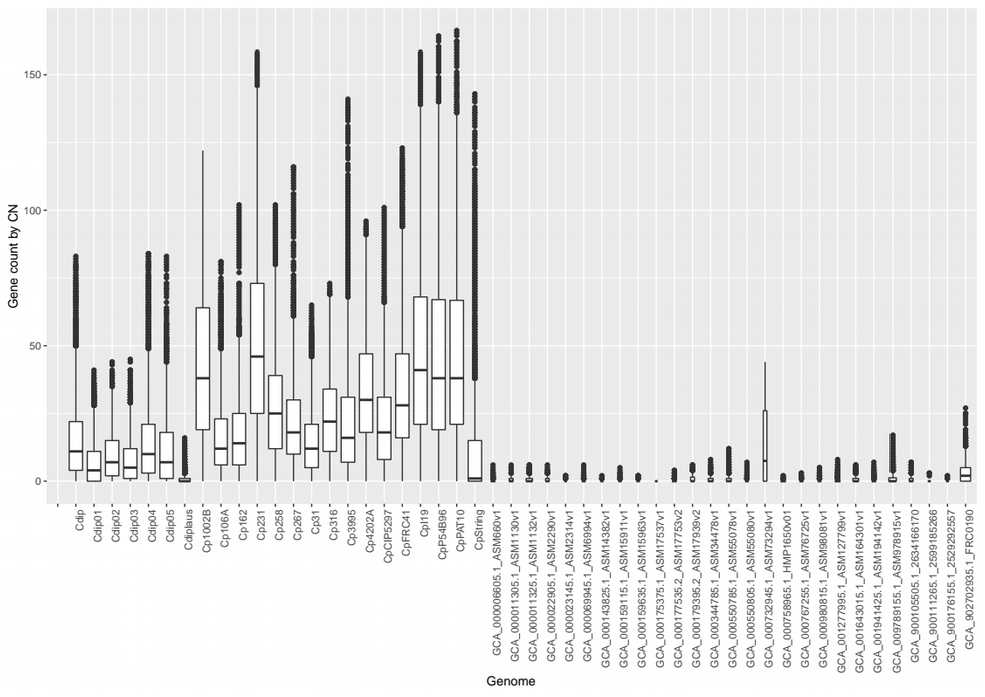
Better possible separation of the genus *Corynebacterium* achieved using the conserved neighbourhood. We achieved splitting via the expansion of a window of pitch three. The stopping criterion was a reduction by more than half in the number of conserved loci for window size. In this query, we did not ensure a full split of *C. pseudotuberculosis* biovars ovis and equi but of the main represented species.

When the GENPPI program runs without the restriction of the threshold window for conserved neighbourhood analysis with progressive increases of -ws until the conservation quality decreases, we call this process a dynamic expansion. In Fig. 6, the measure of genes conserved in a neighbourhood (dynamic extension with -ws 3) showed a high similarity between the genomes of the biovar equi of the species *C. pseudotuberculosis*, strains Cp106A, Cp162, Cp258, Cp31, Cp316, and CpCIP5297. The median of the six equine genomes remained below 25 genes. Within this graph, three out of sixteen genomes of *C. pseudotuberculosis* have box plots with the median below 25 not belonging to the biovar equi, the genomes Cp267, Cp3995, and CpString. We know the genomic relations between the biovars ovis and equi from the literature of *C. pseudotuberculosis* (Soares *et al.*, 2013). When the dynamic expansion step -ws is equal to 1, we have seven out of ten genomes of *C. pseudotuberculosis* biovar ovis whose medians approach those of the biovar equi genomes (data not displayed). However, if we increase the neighbourhood conservation window’s pitch, for example, to -ws 5 and -ws 7, there will be no changes against the result with -ws 3 (data not displayed). Thus, the value that best created the separation of the biovars ovis and equi regarding the gene neighbourhood’s conservation was a dynamic extension step with window size equal to 3, value derived from experimentation and comparison between results.

Nevertheless, in Fig. 6, when we utilized dynamic expansion, the seven genomes of *C. diphtheriae* had medians lower than the lowest median obtained for most *C. pseudotuberculosis* strains. The median of the *C. diphtheriae* species remained lower than the average of most of the species *C. pseudotuberculosis*. The Cdiplaus genome was at a median well below those of the other analysed genomes of *C. diphtheriae*. Considering that the literature reports Cdiplaus as a heterotypic synonym of *Corynebacterium belfantii* (Badell *et al.*, 2020), we have evidence indicating that our analysis of the median genomes of *C. diphtheriae* would provide a correct classification of all genomes of the species *C. diphtheriae* analysed in this study. In addition, the difference between the CpString median compared to all other genomes of *C. pseudotuberculosis* and even with *C. diphtheriae* is noteworthy. In graphs of the number of genes per conserved neighbourhood generated by GENPPI, medians with values close to zero are found for genomes that have only one specimen per species among the analysed set.

The GENPPI’s dynamic expansion to CN makes us pay the price for more accurate mappings. The number of protein comparisons is similar to compound interests; we have a principal multiplied by interest, but a power of ws. The interest is a factor we named ρ and depends on the average number of proteins among the genomes analyzed. To *Corynebacterium*, ρ is equal to 1.15. The result is 2 hours to finish considering a window size equal to three and 50 genomes. However, for *Staphylococcus*, with ρ=1.33 and 57 genomes, we spent 32 hours on the same window pitch. We present a more thorough analysis in Supplementary Method S5. The counterpart of the dynamic expansion algorithm to CN is the fixed retraction. Instead of exponential complexity, we have a logarithmic one, which takes about 40 minutes to process the same 50 *Corynebacterium* genomes, considering an initial window of size 10. Although the dynamic expansion to CN showed the best possible result in distinguishing between species and biovars, we also made analyses with fixed retraction with an initial window size equal to 10 (Supplementary Result Fig. 2).

### 3.4 Interaction networks created with GENPPI for *Corynebacterium*

We submitted a set of 50 genomes of *C. pseudotuberculosis* to several combinations of GENPPI parameters. The analyses were divided between the two types of window sets for a conserved neighbourhood (fixed or dynamic) versus the seven possible types of configurations for a boundary of phylogenetic profiles, including an option that does not restrict the load of interactions mapped in the final report.

It is important to note that the algorithms employed for assessing conserved PP and CN work independently. Each algorithm generates variant sets of interactions that can occur for the same pair of genes. We chose to explore CN execution variations without changing the PP execution mode. The objective was to facilitate the comparison between results. The most relevant results produced by GENPPI, considering the purpose of studying centrality measures, are presented in Table 1. We employed a network generated by STRING software for the genome of *C. pseudo-tuberculosis* as a reference (Ref) for the metrics.

**Table 1.** Metric values obtained for interaction networks by CN and PP.

| Id | CN expansion | Nodes | Medium degree | Density | Edges | Maximum degree |
| --- | --- | --- | --- | --- | --- | --- |
| Ref | - | 2213 | 180.83 | 0.082 | 200088 | 901 |
| f1 | fixed | 2149 | 319.859 | 0.398 | 943553 | 1343 |
| f3 | fixed | 2073 | 287.354 | 0.139 | 297842 | 714 |
| f2 | fixed | 2066 | 70.421 | 0.034 | 72745 | 713 |
| f4 | fixed | 1999 | 132.719 | 0.066 | 132653 | 410 |
| d1 | dynamic | 2150 | 897.189 | 0.417 | 964478 | 1387 |
| d3 | dynamic | 2075 | 317.163 | 0.153 | 329057 | 807 |
| d2 | dynamic | 2027 | 72.109 | 0.036 | 73082 | 706 |
| d4 | dynamic | 1999 | 165.843 | 0.083 | 16576 | 546 |
| d5 | dynamic | 2058 | 272.208 | 0.132 | 280102 | 713 |

Compared to the fixed expansion, the values of the metrics for dynamic expansion in Table 1 were approximate with point exceptions. The highlight is observed because of the average degree of execution with the f1 id. If we exchange only the expansion from fixed to dynamic, we obtain a significant increase in the medium degree’s conservation, with an increase from 320 to 897 being observed. This increase is probably observed because of evolutionary conservation in the genomes of the genus *Corynebacterium*.

The particular web created by d5 Id has a density and average degree above what we consider ideal for the study of centrality measures compared to the STRING reference. However, this network generated the best phylogenetic separation between species via CN (Fig. 6). This result is an example of the flexibility of network generation provided by GENPPI. Our software enables the creation of interaction networks customized for a user’s specific need, such as the study of measures of centrality (lower density) or the study of protein clusters (higher density). Regardless of the end-user objective and considering that interactions have a valid biological meaning, we guarantee the correction of the networks obtained in further studies.

Given the variations in the bulk of vertices and edges that can compose each network created by GENPPI, we expect to experience diversity in the topology of nets created by our software. We present the results of an examination of topology’s variety in Supplementary Result Fig. 3.

## 4 Conclusion

The study of bacterial network topologies based on evolutionarily predicted relationships is a promising area of research. Until this study was conducted, few studies had performed such a query for a genome. A possible cause for this limitation is the absence of software to predict interaction networks from protein sequences alone. Our software presented in this report is a useful tool for any researcher to use. GENPPI can help fill the gap concerning the considerable number of novel genomes assembled monthly and our ability to process interaction networks considering the noncore genes for all completed genome versions. With GENPPI, a user dictates how many and how evolutionarily correlated the genomes are in order to answer a specific scientific query. We offer various configuration modes to employ, ranging from fast and lightweight to more careful and intense computations. However, we should warn users of the usual traps of extensive computational inquiries. Regardless of the chosen processing method, the user can be assured of obtaining a mostly reasonable answer, at least (Esch and Merkl, 2020). We are confident in the GENPPI software because the majority of the necessary relationships that it provided were determined to be correct by CN and PP, as phylogenetic analyses of these relations correctly separated bacterial species. Our software is open-source, and we can compile it for different operational systems.

## Acknowledgements

The authors thank the below-listed funding agencies.

